# Breast cancer brain metastases show increased levels of genomic aberration based homologous recombination deficiency scores relative to their corresponding primary tumors

**DOI:** 10.1101/243535

**Authors:** M. Diossy, L. Reiniger, Zs. Sztupinszki, M. Krzystanek, K. M. Timms, C. Neff, C. Solimeno, D. Pruss, A. C. Eklund, E. Tóth, O. Kiss, O. Rusz, G. Cserni, T. Zombori, B. Székely, I. Csabai, Z. Szallasi

## Abstract

Due to its mechanism of action, PARP inhibitor therapy is expected to benefit mainly tumor cases with homologous recombination deficiency (HRD). Various measures of genomic scarring based HRD scores were developed as a companion diagnostic in order to correlate it with PARP inhibitor sensitivity. We compared a variety of HRD scores in primary tumors and their corresponding brain metastases and found a significant increase in this measure in brain metastases for all measures of HRD that were tested. This discrepancy warrants further investigation to assess whether this observation is common to other metastatic sites, and potentially a significant adjustment of strategy in the application of HRD measures in clinical trials for the prioritization of patients for PARP inhibitor therapy.

**Key message:** We quantified homologous recombination deficiency (HRD) in paired primary breast cancer and brain metastases samples based on a previously published data set. We showed that HRD significantly increases in the brain metastases relative to its paired primary tumor. An independent validation was also performed on another set of primary breast cancer and brain metastasis pairs using the “my choice” HRD score in collaboration with Myriad Genetics.

These confirmatory results suggest that brain metastases of breast cancer tend to have significantly higher HRD scores, which would prioritize those patients for PARP inhibitor therapy with those agents that cross the blood brain barrier such as veliparib and niraparib.

## Introduction

As a result of recent developments in treatment regimens, survival in patients with metastatic breast cancer has improved. However, the benefit of improved survival is mitigated by the fact that 15–20 % of patients with metastatic breast cancer will develop brain metastases[1]. Current therapeutic approaches have not lowered this incidence, on the contrary, probably due to longer survival, the incidence of brain metastasis in breast cancer patients is rising[1]. Brain metastases have been associated with poor prognosis and with neurological impairments severely affecting quality of life[2]. Due to its unique location, and perhaps its distinct biology, treatment options in brain metastasis have been limited.

Homologous recombination (HR) is an error-free mechanism of double-stranded DNA breaks repair. HR is often deficient in breast and ovarian cancer sensitizing them to PARP inhibitor or platinum-based therapy[3–6]. Due to its remarkable clinical efficacy the first PARP inhibitor, olaparib, received accelerated approval by the Food and Drug Administration (FDA) for BRCA mutant ovarian cancer treatment. Two other PARP inhibitors (niraparib and rucaparib) also received FDA approval recently and others, such as veliparib, are also in clinical development. Other cancer types associated with BRCA mutations, such as breast and prostate cancer, also show promise in PARP inhibitor trials[6–8]. More recently, olaparib monotherapy showed significant benefit over standard therapy in HER2-negative, germline BRCA1/2 mutant metastatic breast cancer patients[6].

Considering these promising clinical results, the question arises whether a similar significant clinical benefit can be achieved with PARP inhibitors in the case of brain metastases as well.

To achieve such therapeutic benefit at least two criteria need to be satisfied. Firstly, the agent should be able to cross the blood brain barrier in order to reach the site of therapeutic action. This may hold true for some of the PARP inhibitors because both niraparib[9] and veliparib[10] were shown to cross the blood brain barrier, making those at least potential candidates for the treatment of brain metastases.

Secondly, the brain metastases should present HR deficiency, to be sensitive to this type of treatment.

Considering the mechanistic basis for its action (synthetic lethality with HR deficiency), usually three groups of patients are evaluated for their response to PARP inhibitors. Germline BRCA1/2 mutant patients (with perhaps the addition of mutation carriers in other HR genes, such as Rad51C, PALB2 etc.) get the most significant benefit from PARP inhibitor treatment. Only a minority (10% or less) of breast cancer brain metastasis patients will fall into this category. While BRCA1/2 mutation is an important determinant of HR deficiency, tumors without BRCA mutations can also be HR-deficient, and therefore, also be sensitive to PARP inhibitor-based therapy. Consequently, the BRCA wild type cases are evaluated separately according to their HR deficiency status. Such BRCA1/2 wild type cancer cases with likely HR deficiency tend to show less, but still significant, benefit from PARP inhibitor treatment relative to BRCA mutation carriers. Conversely, BRCA1/2 wild type cancer cases without HR deficiency showed the least, rather marginal benefit from PARP inhibitor treatment emphasizing the role of HR deficiency in PARP inhibitor sensitivity[4].

Determining HR deficiency in a BRCA1/2 wild type background, which is essential for the above-described clinical evaluation, has not been a trivial task. We developed a DNA “scarring” signature, telomeric allelic imbalance (tAI) that detects and quantifies the level of homologous recombination deficiency in human tumor biopsies[11]. This method was later combined with two other methods and it is currently part of the myChoice HRD test[3]. Recently, with the availability of a significant amount of whole genome sequencing data on breast cancer, another genomic aberration-based HR deficiency measure was developed, by adding single nucleotide variation and large scale genomic rearrangements based signatures to genomic instability analysis[12].

So far, it has not been determined how often breast cancer brain metastases show HR deficiency. This was the main goal of this project. We have calculated HR deficiency measures in previously published next generation sequencing data of primary breast cancer/brain metastasis pairs[13]. The initial analysis was followed up by a direct measurement of the myChoice HRD test in an independent cohort.

## Results

We calculated the three previously published homologous recombination deficiency associated “genomic scar” scores, which were originally developed using whole genome SNP arrays on previously published whole exome next generation sequencing data obtained from paired primary breast cancer and brain metastases biopsies. We found that all three WES-based scores significantly increased in the brain metastases. (Figure 1) Recently, in addition to the above mentioned three scores that were originally based on measurements with SNP arrays, a different measure of HR deficiency, “HRDetect”, was published that incorporated single nucleotide variations, short indels and large scale genomic rearrangements as well[12]. “HRDetect” was developed using WGS data and it is not directly transferable to whole exome sequencing data. Since the dataset of brain metastasis/primary tumor pairs contained only WES data, we extracted the features of HRDetect that could be calculated based on the available sequences and retrained HRDetect on those. Using the same principles as the original HRDetect publication, we reconstructed a similar complex measure of HR deficiency using only WES data. (Details of converting the WGS based HRDetect measure into a WES based HRDtetect measure are available in the supplementary material, section 4).

We found that this newly reconstructed, “WES-HRDetect” model also distinguished well between HR-deficient and HR-proficient primary breast cancer, as determined by e.g. BRCA mutation status (supplementary material, section 4). Therefore, we calculated the WES-HRDetect scores for the primary breast cancer brain metastasis pairs as well.

**Figure 1:**
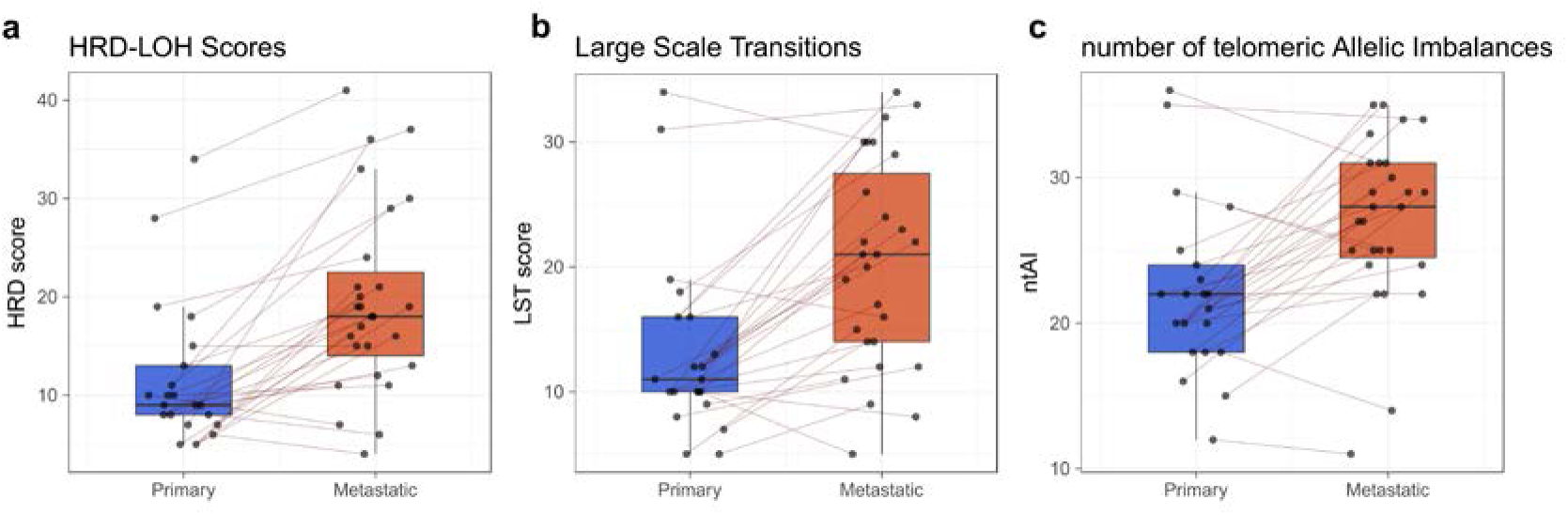
Genomic scar scores of the breast cancer brain metastasis samples. Distributions of the HRD-LOH (a), LST (b) and ntAI (c) scores. The corresponding primary-metastatic pairs are connected by thin red lines. Scores for each of the measures were increased in metastases compared to primary tumors with p-values of the paired t-tests: pLOH= 5.13e-6, pLST= 1.12e-6 and pntAI= 1.63e-5 respectively.

Compared to the results obtained by the HRD scores, this new measure showed an even more substantial increase of HR-deficiency in brain metastases relative to the primary tumors (Figure 2).

**Figure 2:**
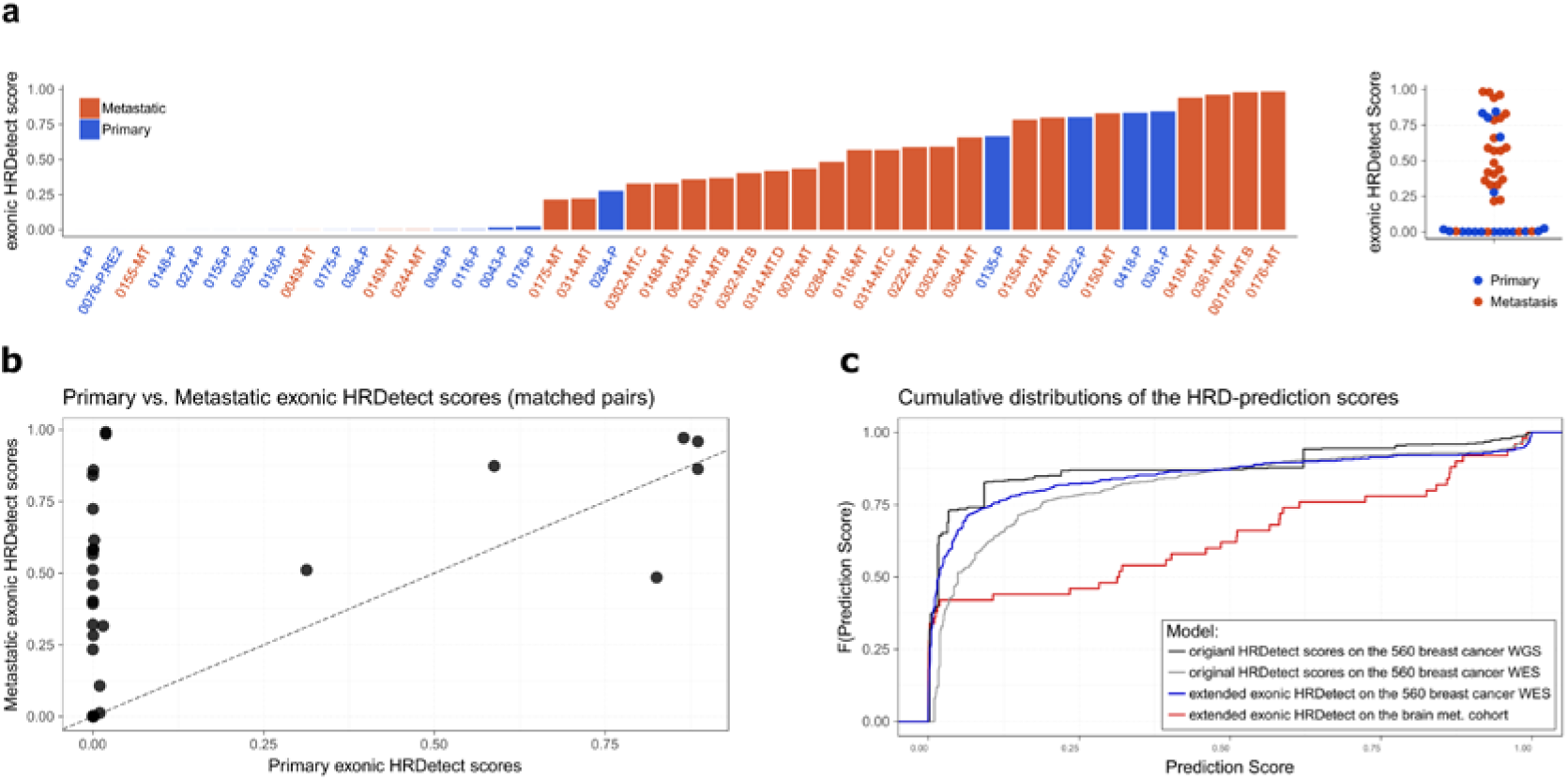
Exonic HRDetect scores. a) Extended HRDetect scores (HRD-LOH scores included) of the primary and metastatic samples. b) Primary HRDetect scores vs. Metastatic HRDetect scores. Noticeably, there is only a single case (PB0222) for which the primary sample’s score is higher than its metastatic pair’s. c) Cumulative distributions of the original HRDetect models (WGS and WES) and the extended model’s scores on the 560 breast cancer samples and on the breast cancer brain metastasis cohort.

We sought independent validation of these observations by assembling a cohort of primary breast cancer brain metastasis pairs, which were processed for the myChoice HRD score as described in[3]. Figure 3 shows that in the majority of the cases the brain metastasis showed a higher myChoice HRD score than the primary tumor. Furthermore, in five cases the brain metastasis “switched myChoice HRD status” relative to the primary tumor from a low myChoice HRD score (HRD-) to a high myChoice HRD score (HRD+), which is based on the currently accepted threshold value of 42[3].

**Figure 3:**
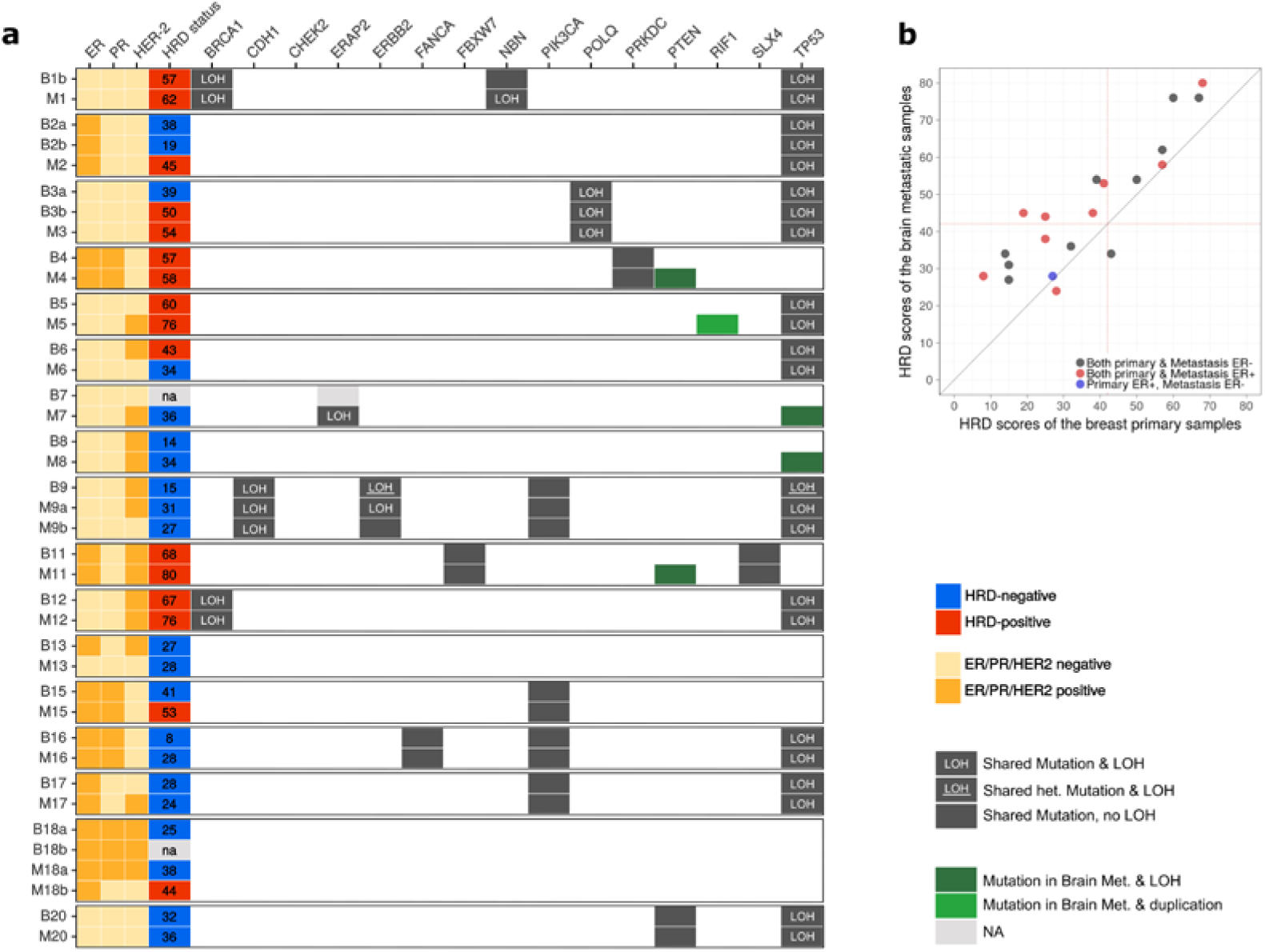
Summary of the control samples. a) Hormone receptor and HER-2 status along with the mutational profile and HRD status of the control samples determined by Myriad Genetics. In the majority of cases (14/17) HRD score was higher in the metastasis compared to the primary tumor. In 3 cases, all of which was ER positive, the HRD status of the metastasis switched to positive from an HRD negative primary. Mutations of either PTEN, RIF1 or TP53 were gained in 5 of the brain metastases. LOH was more common in the brain metastases. b) Genomic scar scores determined by Myriad Genetics summed together into a single HRD-score. HRD scores were increased in metastases compared to primary tumors (p = 5.4e-6). The figure is divided into 4 quadrants by the HRD-score = 42 lines, among which the top left quadrant contains the therapeutically relevant cases. Two of these samples (marked with asterisks on the plot) belong to the same patient (patient 2).

Considering the significant changes found in all of the HR deficiency scores studied between primary tumors and brain metastases, we also investigated whether further differences can be found in other mutational signatures as well, such as those published by Alexandrov et al.[14] (Figure 4). We found that, as it is already well known in the field, primary breast tumor mutational profiles are dominated by signature 1 (aging related signature), with lesser contributions from signature 2 (APOBEC signature), signature 3 (BRCA mutation associated), signature 5 and signature 6. Strikingly, signature 3, the BRCA mutation associated signature significantly increased in about one third of the brain metastatic cases, which is in accordance with the findings above, since this signature is part of the HRDetect score.

**Figure 4:**
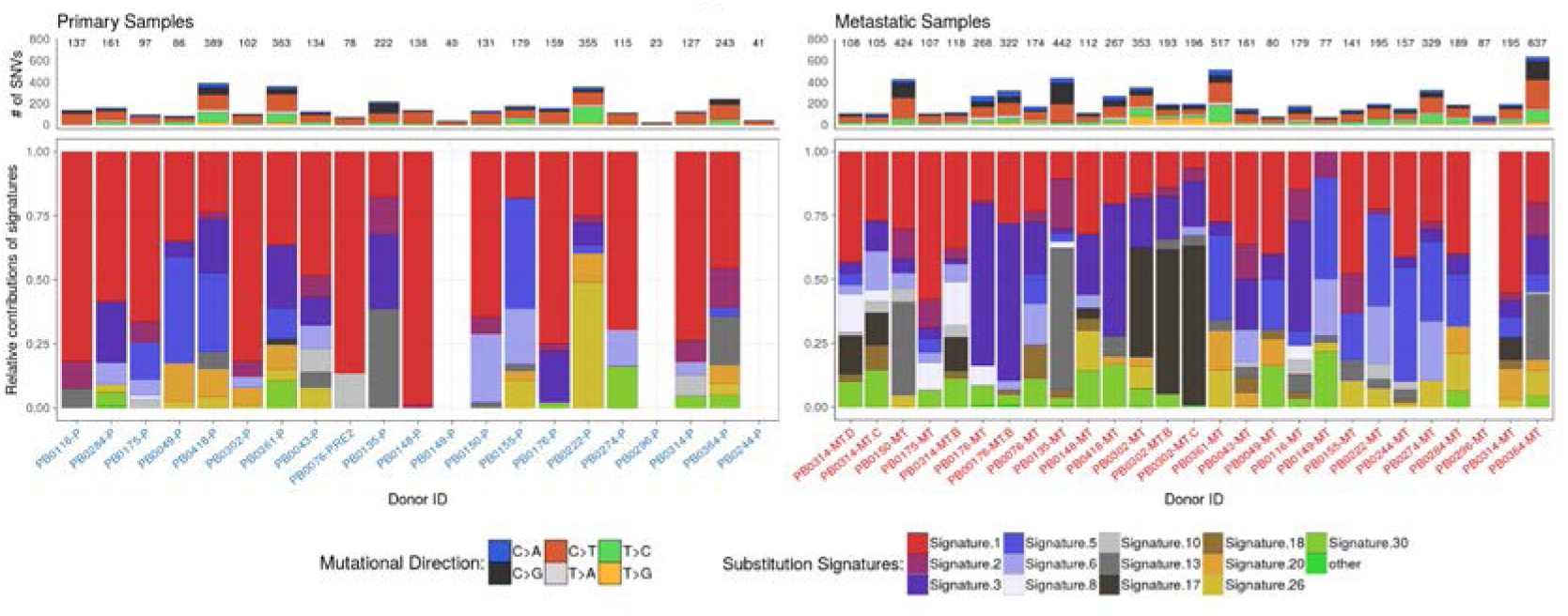
Summary of somatic point mutations. Top panels: Total number of point mutations in the samples colored by the relative abundance of the 6 mutational directions. Bottom panel: Shares of the somatic point-mutation signatures (REF here) in the triplet mutational profiles of the samples. The reconstruction errors (REF here) of specimens 0149-P, 0296-P, 0244-P and 0296-MT did not pass the cosine similarity threshold, hence those samples have no bars on the plots.

Interestingly, in about 20–25% of the brain metastasis cases signature 5, (another aging related signature distinct from signature 1), has become similarly dominant as signature 1. Finally, in a single case (0302), signature 17 (unknown relevance and biological origin) became the predominant mutational signature in the breast cancer brain metastasis. We also investigated whether the changes in HR deficiency measures is correlated with the time elapsing between the detection of the primary tumor and the brain metastasis and found significant correlation in the case of both cohorts (Supplementary Figure 5.1).

## Discussion

This work is the first demonstrating that the various DNA aberration based HRD measures are significantly higher in brain metastases relative to their primary breast tumor counterparts. The possibility of such an increase was previously suggested in a publication showing that a BRCA1 deficient-like gene expression signature is higher in breast cancer brain metastases, but that study did not use patient matched primary metastasis sample pairs[15].

This discrepancy has several clinical consequences. Ideally, only patients sensitive to a given therapy should receive that particular treatment. In the case of PARP inhibitors, it is likely that mainly patients with HR deficiency benefit from this form of therapy. Therefore, correlating the clinical benefit of PARP inhibitor therapy with HR status of the tumors has been intensely investigated[4]. It should be noted, however, that HR status is often determined in biopsies derived from the primary tumor but clinical benefit is determined at a more advanced, often metastatic stage of the disease. If the HR deficiency status is significantly different in the metastases relative to the primary tumors then the observed correlations will be corrupted leading to contradictory clinical results. For example, in the case of breast cancer repeated observations demonstrated a correlation between the ntAI score and response to platinum based therapy[11]. The TNT trial[16], however, studied the efficacy of platinum based therapy in locally advanced or metastatic triple-negative breast cancer and failed to find a correlation between the myChoice HRD score and response to carboplatin. It should be noted, however, that the myChoice HRD score was determined on the primary tumors and not on the metastases. The discrepancy between the myChoice HRD score of the primary and metastatic sites presented in this study may be partially responsible for this lack of correlation.

There are several ongoing clinical trials evaluating the potential clinical benefit of PARP inhibitor treatment in metastatic breast cancer, including NCT02723864 (https://clinicaltrials.gov/ct2/show/NCT02723864). This trial, however, excludes patients with active brain metastasis. Based on our results, it seems likely that breast cancer brain metastases might be a particularly sensitive population. The SWOG trial S1416 is a Phase II Randomized Placebo-Controlled Trial of Cisplatin with or without ABT-888 (Veliparib) in Metastatic Triple-Negative Breast Cancer (TNBC) and/or BRCA Mutation-Associated Breast Cancer. This trial includes patients with brain metastasis but excludes patients with estrogen receptor positive breast cancer. As we showed in our validation cohort, the majority of estrogen receptor positive breast cancer brain metastasis cases showed high myChoice HRD scores. It might be worth establishing the proportion of HRD+ ER+ breast cancer cases on a larger cohort as those patients might be responding to PARP inhibitors as well.

The significant increase in HR deficiency scores in brain metastases may arise due to at least two distinct mechanisms. 1) Clonal selection of tumor cells with higher HR deficiency score. Tumors cells with higher levels of genomic instability, higher levels of HR deficiency may be able to adjust more readily to a distinct environment such as the brain. 2) It is also possible that the growing brain metastases becomes gradually more genomically unstable due to some unidentified mechanism. Both mechanisms are consistent with the fact that the time elapsing between primary and brain metastases often takes up to 5–10 years in breast cancer, and the length of this time showed a strong correlation with the increase in the various HRD measures in brain metastases.

Brain metastases of breast cancer patients are surgically removed with palliative intent in a significant number of cases. According to the guidelines, surgical resection is recommended in patients with a limited number (1 to 3) of newly diagnosed brain metastases, especially with controlled systemic disease and good performance status[17]. In such cases, when the histological material is already available, it might be worth considering determining the HR deficiency score in order to determine the likely sensitivity of the tumor to PARP inhibitors versus other second-/third-line therapeutic options.

## Materials and methods

### Whole exome samples

The whole exome sequencing (WES) data of the matched germline-primary-metastatic trios, under accession number phs000730.v1.p1, was downloaded from the database of Genotypes and Phenotypes (dbGaP)[13]. The dataset contains binary alignment (BAM) files of primary tumors and their corresponding metastatic pairs from 86 patients. Twenty-one of the primary sites were located in the breast, and all primary tumors had at least one matched intracranial metastasis. This breast cancer cohort formed the basis of our initial investigations.

### Independent validation cohort

We collected FFPE tissues from 17 cases with primary breast cancer and corresponding brain metastases. Permissions to use the archived tissue have been obtained from the Regional Ethical Committee (No: 510/2013, 86/2015) and the study was conducted in accordance with the Declaration of Helsinki.

### Somatic Mutation Calling

The samples were pre-processed, aligned to the hg19 reference genome with the BWA mem algorithm, and post-processed (base-recalibration) by the original authors[13]. The good quality of the reads in the binary alignment files was confirmed using fastQC (http://www.bioinformatics.babraham.ac.uk/projects/fastqc/). We called somatic substitutions along with short insertions and deletions with MuTect2, a part of the Genome Analysis Toolkit (GATK 3.7)[18]. The majority of the samples were derived from formalin fixed, paraffin embedded specimens (FFPE), which are well known to contain mutational biases (mostly C>T/G>A transitions), due to an enhanced deamination rate of cytosines caused by formalin)[19]. We used an approach analogous to the OxoG filter[20] to remove mutations that were consistent with this FFPE-bias (details are available in the supplementary material, section 1).

### Classification of deletions

Deletions were grouped into three classes: 1) if the entire deleted sequence of a single event was repeated after the breakpoint, it was considered a repeat, 2) if only the first n nucleotides were repeated after the breakpoint, it was considered a size n microhomology, and 3) if it had no such characteristics, it was treated as a unique deletion. In order to eliminate those microhomologies that have appeared by pure chance instead as the result of the activity of the microhomology mediated end joining DNA-repair pathway, they had been further divided into n < 3 and n >= 3 subgroups. In our prediction models only the n >= 3 subgroup was considered.

### Copy Number Analysis

Deriving copy number data from the whole exome sequences was executed using the sequenza R package[21]. The ranges of ploidy and cellularity were confined into the [1,5] and [0.1,1] intervals respectively, and segmentation data was generated for autosomes only. The final solutions were inspected manually and if an alternative solution provided a more reliable ploidy and cellularity estimate than the default one (with the highest log-posterior probability), the alternative model was selected instead.

### Calculation of Genomic Scar scores

The three genomic scar scores were calculated from the segmentation data returned by sequenza, using their standard definitions[11, 22, 23]. In brief, the number of telomeric allelic imbalances (tAI) is the number of subtelomeric regions that exhibit allelic imbalance that start beyond the centromere and extend to the telomere, large-scale state transitions (LST) count the number of chromosomal breaks between adjacent regions of at least 10 Mb, and the loss of heterozygosity score (HRD-LOH) measures the number of regions with LOH which are larger than 15 Mb, but shorter than the whole chromosome. (Further information is available in the supplementary material, section 3.)

### Statistical considerations

For statistical comparison we have formed primary-metastatic pairs from the samples. Out of the 21 primary specimens 3 had multiple metastases which were obtained from distinct intracranial tumors. Such metastases were compared to the same primary source. This choice was more reasonable than taking the average properties of multiple metastatic samples, since they might have originated from different primary colonies, and might have gone through different micro-evolutionary paths (Supplementary Table 1). The average scores of the three genomic scars were then compared between the primary and metastatic samples by using paired sample t-tests with two tailed alternative hypotheses.

### Extraction of mutational signatures

Mutational catalog and signature extractions were performed by the deconstructSigs R package[24]. For mutational extraction only those samples were considered, which had at least fifty somatic substitutions and the assessment of signature contributions was restricted to those signatures that were previously reported to be operative in breast cancers (Signatures 1, 2, 3, 5, 6, 8, 10, 13, 17, 18, 20, 26 and 30)[14]. To verify the computational process, the original spectra were reconstructed from the extracted signatures, and the cosine similarity between the two 96-dimensional vectors was calculated. If this similarity was smaller than 0.8, the extraction was considered unsuccessful and the affected samples were excluded from any further analysis.

### The alternative exonic HRDetect model

The original HRDetect model [12] was trained on mutational features derived from 560 breast cancer whole genome sequences, and although the authors have created an artificial whole exome variant (limiting WGS to typical WES genomic coverage area), the exact details of this computational pipeline were not published. Therefore, in order to use this predictor for our WES samples, we had to retrain the LASSO logistic regression model on the 560 artificial WES dataset, with the HRD indices included[12]).

Our intentions were to carefully replicate all the steps of the original article. We used the same data transformations and the tenfold nested cross-validation strategy to assess the generalizability and robustness of the final weights (details available in the supplementary material, section 4). The final weights (Supplementary Table 7.) were acquired with λ =0.00543. The most significant weight in this model was the HRD index, and its inclusion has increased the sensitivity of the predictor model from 0.727 (sensitivity of the original model trained on exomes) to 0.766 (Supplementary Figure 4.4).

### myChoice HRD Analysis

DNA was extracted from FFPE tissue sections and analysis performed as previously described [3]. Briefly, genome wide SNP data and sequence spanning the coding regions of 111 genes was generated using a custom hybridization panel. Details of the methods used for identification of sequence variants are provided in[3]. Allelic imbalance profiles were generated to determine the scores for 3 individual biomarkers of genomic instability (TAI, LOH, and LST). The combined HRD score is the sum of the 3 independent biomarkers. A myChoice HRD threshold of 42 has previously been developed to identify HR deficient tumors using this test[3]. Tumors are considered HRD+ if they have a high myChoice HRD score (≥42) or a tumor BRCA1/2 mutation and HRD-if they have a low myChoice HRD score (<42) and wild-type BRCA1/2.

## Acknowledgment

This work was supported by the Research and Technology Innovation Fund (KTIA NAP 13-2014-0021 to Z.S.); Breast Cancer Research Foundation and the Novo Nordisk Foundation Interdisciplinary Synergy Programme Grant (NNF15OC0016584 to Z.S.), by the ÚNKP-17-4-III-SE-63 New National Excellence Program of the Ministry of Human Capacities to L.R. and by Tesaro Inc.

## Online Supplementary material

**supplementary_notes.pdf**: A summary about the steps, that were followed during the analysis. It contains 16 supplementary figures, numbered according to the chapter number of the text, and 5 small supplementary tables numbered from 5 to 9. Supplementary tables 1–4 are available in separate excel sheets:

**Supplementary Table 1:** The reanalyzed dataset of paired breast primary brain metastatic pairs, and their genomic scar scores

**Supplementary Table 2:** Mutational profile of the validation breast cancer brain metastasis cohort.

**Supplementary Table 3:** a) Detailed analysis of the validation cohort by Myriad Genetics, b) Genomic scar scores of the validation samples.

**Supplementary Table 4:** Exonic HRDetect predictors of the reanalyzed dataset.

